# Aether: Leveraging Linear Programming for Optimal Cloud Computing In Genomics

**DOI:** 10.1101/162883

**Authors:** Jacob M. Luber, Braden T. Tierney, Evan M. Cofer, Chirag J. Patel, Aleksandar D. Kostic

## Abstract

Across biology we are seeing rapid developments in scale of data production without a corresponding increase in data analysis capabilities. Here, we present Aether (http://aether.kosticlab.org), an intuitive, easy-to-use, cost-effective, and scalable framework that uses linear programming (LP) to optimally bid on and deploy combinations of underutilized cloud computing resources. Our approach simultaneously minimizes the cost of data analysis while maximizing its efficiency and speed. As a test, we used Aether to *de novo* assemble 1572 metagenomic samples, a task it completed in merely 13 hours with cost savings of approximately 80% relative to comparable methods.

## Main

Data accumulation is exceeding Moore’s law, which only still progresses due to advances in parallel chip architecture.^1^ Fortunately, the shift away from in-house computing clusters to cloud infrastructure has yielded approaches to computational challenges in biology that both make science more reproducible and eliminate time lost in high-performance computing queues^2,3^; however, existing off-the-shelf tools built for cloud computing often remain inaccessible, cumbersome, and in some instances, costly.

Solutions to parallelizable compute problems in computational biology are increasingly necessary; however, batch job-oriented cloud computing systems, such as Amazon Web Services (AWS) Batch, Google preemptible Virtual Machines (VMs), Apache Spark, and MapReduce implementations are either closed source, restrictively licensed, or locked in their own ecosystems making them inaccessible to many bioinformatics labs.^4,5^ Other approaches for bidding on cloud resources exist, but they neither provide implementations nor interface with a distributed batch job processing backend.^6–8^

Our proposed tool, Aether, leverages a linear programming approach to minimize cloud compute cost while being constrained by user needs and cloud capacity, which are parameterized by the number of cores, RAM, and in-node solid-state drive space. Specifically, certain types of instances are allocated to large web service providers (e.g., Netflix) and auctioned on a secondary market when they are not fully utilized.^6^ Users bid amongst each other for use of this already purchased but unused compute time at extremely low rates (up to 90% off the listed price).^9^ However, this market is not without its complexities. For instance, significant price fluctuations, up to an order of magnitude, could lead to early termination of multi-hour compute jobs (Figure 1A). Clearly, bidding strategies must be dynamic to overcome such hurdles.

**Figure 1.**
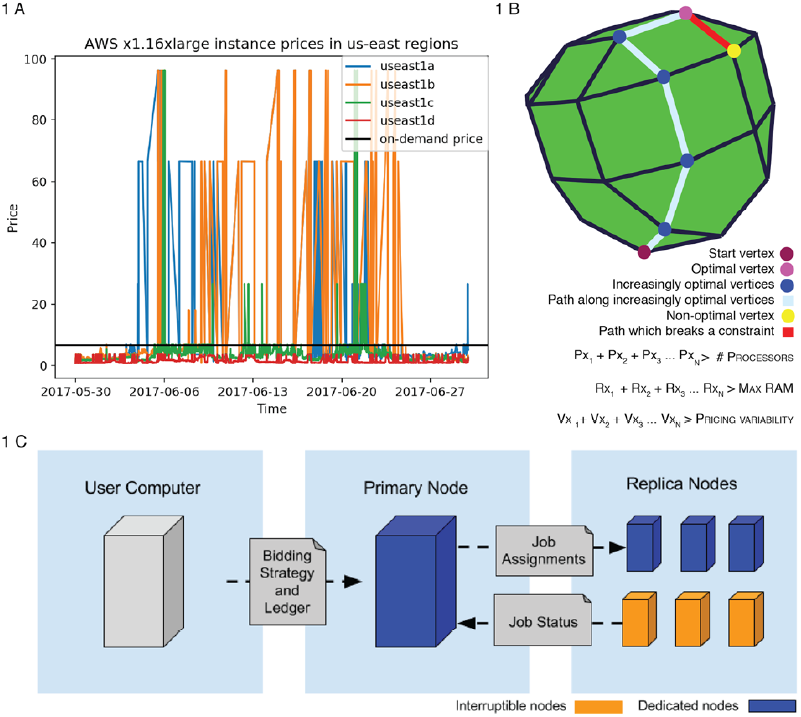
A) Pricing history of an x1.16xlarge EC2 Instance on AWS showcasing variability of an order of magnitude, in both directions, for spot prices. B) Simplified example showing three constraints on a sample bidding approach minimizing an objective function cTx considering cost according to a system of constraints represented as inequalities. x1, x2, and x3 represent the number of specific types of compute nodes to solve for. Each inequality represents a constraint and adds another dimension to the space which the simplex algorithm needs to traverse vertices in to find ideal solution. The actual system of inequalities has 140 constraints. The green line represents the optimal solution. C) An overview of Aether’s computing infrastructure.

Due to this pricing variability, it can be optimal to bid on non-auctioned instances in certain regions. To properly handle this case, we include additional linear constraints for both an instance’s on-demand and at-auction prices. The solution vector is bounded by the number of currently running instances as well as limits due to provider capacity. Finally, to avoid bidding on instances that will spike in price, the algorithm looks at pricing history and sets a final constraint corresponding to a user’s maximum tolerable pricing variability. For each run of the bidder, this system of 140 inequalities is converted to slack (standard) form and then solved with the simplex algorithm as implemented in Python’s *scipy.linprog* library (Figure 1B).^10^ This naively outputs suggested compute bids as floats; obviously, a fraction of an instance is not a valid bid and generating integer solutions to linear programming problems is NP-hard. However, a true integer linear programming solution is not required, as the constraints still hold if the floor is taken from each bid, provided that preprocessing is done to remove underutilized instance types and those that cannot process a unary compute job. To reach this optimal integer pseudo-solution, the linear programming solver is run recursively such that these non-feasible fractional bids are iteratively removed. Additionally, adhering to the pricing variability constraint is not guaranteed to yield the optimal value, so the simplex algorithm is applied iteratively, setting the pricing variability from zero to the maximum specified value until either the optimal value is found or it is determined that there is no solution to the system. In the event of finding no solution, the user must re-run the program with a higher maximum cost. This approach results in a tractable average case runtime, which yields essentially instant bidding suggestions given the small size of the system being solved.

Aether consists of bidder and batch job processing command line tools which query instance metadata from the vendor application programming interface (APIs) to formulate the linear programming problem. Subsequently, the replica nodes specified by the linear programming result are launched and placed under the control of a primary node, which assigns batch processing jobs over Transmission Control Protocol (TCP), monitors for any failures, gathers all logs, sends all results to a specified cloud storage location, and terminates all compute nodes once processing is complete (Figure 1C). Additionally, Aether is able to distribute compute across multiple cloud providers. Our implementation runs on any Unix-like system; we ran our pipeline and cost analysis using AWS but have provided code to spin up compute nodes on either Microsoft Azure or on a user’s local physical clusters.

To test our bidding approach and batch job pipeline at scale, we used our framework to *de novo* assemble and annotate 1572 metagenomic, longitudinal samples from the stool of 222 infants in Northern Europe (Figure 2A).^11–14^ The sequencing data within datasets from the DIABIMMUNE consortium ranged from 4,680 to 22,435,430 reads/sample with a median of 19,020,036 reads/sample. Assemblies were performed with MEGAHIT and annotations were done with PROKKA.^15,16^

**Figure 2.**
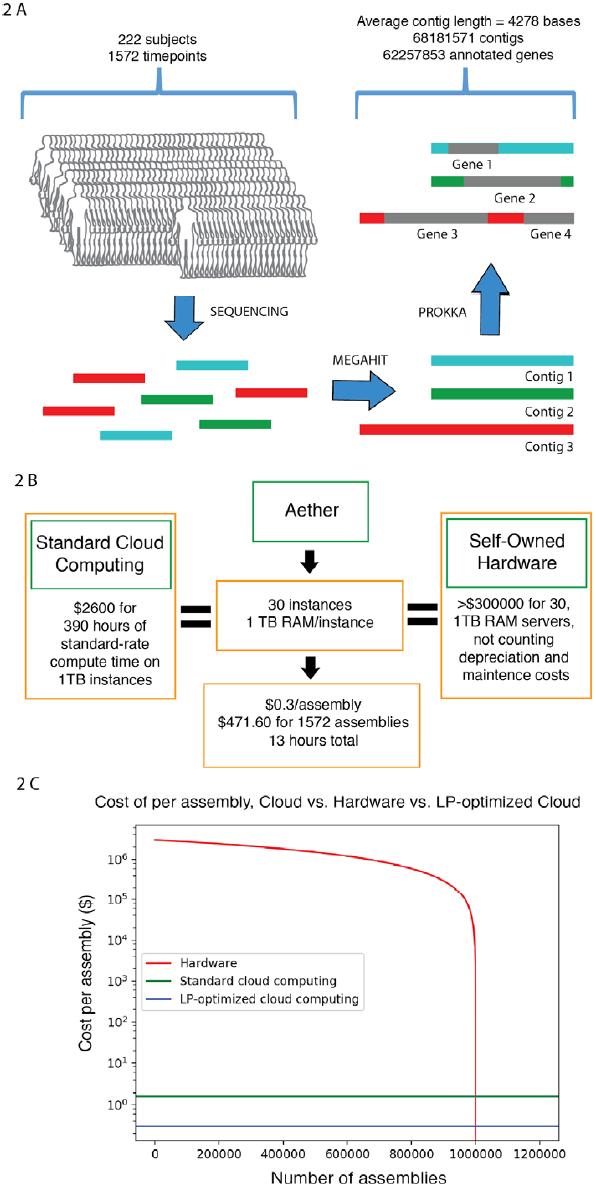
A) Overview of the assembly process. A total of 1572 fecal samples were collected and sequenced at various timepoints during the first 3 years of 222 individuals lives. These were assembled with MEGAHIT into 68,181,571 contigs. Across all samples, a total of 62,257,853 genes, ~1,000,000 of which were unique, were then annotated using Prokka. Only contigs that were over 1000 bases long were used. The mean length of this group was 4278 bases. B) Cost comparison between Aether, standard cloud computing, and user-maintained hardware. Total assembly cost was 18% ($471.60) of what it would have been using on-demand instances. We estimated the upfront cost of a server equivalent to those used to analyze the data being ~$10,000. Given that we used 30 instances of these servers, the total cost of hardware would be $300,000 according to pricing information from Penguin Computing and Dell, not counting system maintenance and depreciation. C) Comparison of cost per assembly for the three platforms outlined in 2B.

Metagenomic data, shotgun DNA sequencing of microbial communities, is difficult to analyze because of the enormous amounts of compute required to naively assemble short sequence reads into large contiguous spans (contigs) of DNA. To accomplish our assemblies, our bidding algorithm suggested that the optimal strategy would be to spin up 30TB of RAM across underutilized compute nodes. Our networked batch job processing module utilized these nodes for 13 hours and yielded an assembly and annotation cost of ~$0.30 per sample (Figure 2B). Theoretically, the pipeline can complete in the time it takes for the longest sub-process (i.e. assembly in this case) to finish (~7 hours). Spinning up the same nodes for this long without a bidding approach would cost ~$1.60 per sample (Figure 2C). In order for on-site hardware to achieve the same cost efficiency as our pipeline, one would have to carry out on the order of 1 million assemblies over the lifespan of the servers, a practically insurmountable task (Figure 2B). Such efficiency in both time and cost at scale is unprecedented. In fact, due to resource paucity, computational costs have forced the field of metagenomics to rely on algorithmic approaches that utilize mapping back to reference genomes rather than *de novo* methods.^17^

In simulated runs of the bidder incorporating pricing history from periods where ask prices were approximately an order of magnitude higher than normal on the east coast of the United States (Figure 1A), Aether suggested utilization of different instance types that would have resulted in similar cost and time to completion as our actual run. To allow users make optimal usage of these benefits, the ability to simulate bidding for different timeframes is included as a feature. By not having to potentially re-run analysis pipelines (due to being outbid on compute during runtime), we claim that utilizing Aether leads to a reduction of market inefficiencies. Future directions include training the bidding algorithm to predict its own effect on pricing variability when being utilized at massive scale as well as distributing compute nodes across datacenters when enough resources are being spun up to strongly influence the market.

To our knowledge, this is the first implementation of a bidding algorithm for cloud compute resources that is tied both to an easy-to-use front-end as well as a distributed backend that allows for spinning up purchased compute nodes across multiple providers. Conceivably, this tool can be applied to any number of disciplines, bringing cost-effective cloud computing into the hands of scientists in fields beyond biology.

## Methods

### Implementation Details

Computational resources and monetary costs are mapped to each available instance type at run-time by querying the cloud providers’ web-based public APIs. To identify the ideal resource selection, we feed these data, along with constraints provided by the user, into our multi-objective optimization procedure. The user-defined set of jobs is subdivided into computational workloads according to the resources available to each node, and distributed across the worker nodes by a central server. In a single node’s workload, jobs are executed in parallel but may complete asynchronously. Upon completion of a job, the replica node notifies the central server, which then schedules another task for the replica. To prevent scheduling errors, we synchronized changes in the primary node’s job ledger, and used at-least-once message delivery. We controlled access to computational resources and accounts with AWS Identity & Access Management (IAM) security groups and Azure Identity and Access Management (IaAM), which their respective providers recommend for authentication and authorization. Additional details regarding Aether’s implementation are available on the project website (http://aether.kosticlab.org).

### Data Availability

Infant metagenomic data utilized are freely available at https://pubs.broadinstitute.org/diabimmune and with EBI SRA accession ERP005989.

### Code Availability

The source code for Aether is available on GitHub (https://github.com/kosticlab/aether), where it is being actively maintained and updated. The examples included in this manuscript, along with documentation and step-by-step tutorials, are available on the project website (http://aether.kosticlab.org).

## Acknowledgements

This work was funded by NIH/NHGRI T32 HG002295, PI: Park, Peter J (J.M.L.), an AWS Research Credits for Education Grant (J.M.L. and A.D.K.), a Microsoft Azure for Research Grant (B.T.T. and C.J.P.), NIH NIEHS R00 ES023504 (C.J.P.), NIEHS R21 ES025052 (C.J.P.), NSF Big Data Spoke grant (C.J.P.), a Smith Family Foundation Award for Excellence in Biomedical Research (A.D.K.), and an ADA Pathway to Stop Diabetes Initiator Award (A.D.K.). We thank Thomas Lane and Chengwei Luo for their feedback and review of the manuscript. We thank Tommi Vatanen for helping us with data access.

## Author Contributions

J.M.L. and B.T.T. conceived the project. J.M.L. and B.T.T. designed and implemented the LP bidding approach. J.M.L, B.T.T, and E.M.C. implemented the distributed batch processing pipeline with guidance from C.J.P. and A.D.K.. C.J.P. and A.D.K. supervised the project. E.M.C. wrote the documentation, tutorial, and other online resources. J.M.L., E.M.C., and B.T.T. wrote the manuscript.

## Competing Interests

The authors declare no competing interests.

## Correspondence

Correspondence and requests for materials should be addressed to C.J.P. or A.D.K.

